# Tissue Level Profiling of SARS-CoV-2 antivirals in mice to predict their effects: comparing Remdesivir’s active metabolite GS-441 524 vs. the clinically failed Hydroxychloroquine

**DOI:** 10.1101/2020.09.16.299537

**Authors:** Oliver Scherf-Clavel, Edith Kaczmarek, Martina Kinzig, Bettina Friedl, Malte Feja, Rainer Höhl, Roland Nau, Ulrike Holzgrabe, Manuela Gernert, Franziska Richter, Fritz Sörgel

**Author notes:** Corresponding author at: IBMP – Institute for Biomedical and Pharmaceutical Research, Paul-Ehrlich-Straße 19, D-90562 Nürnberg-Heroldsberg, Germany.

## Abstract

**Background and Objectives:** Remdesivir and hydroxychloroquine are or were among the most promising therapeutic options to tackle the current SARS-CoV-2 pandemic. Besides the use of the prodrug remdesivir itself, the direct administration of GS-441 524, the resulting main metabolite of remdesivir, could be advantageous and even more effective. All substances were not originally developed for the treatment of COVID-19 and especially for GS-441 524 little is known about its pharmacokinetic and physical-chemical properties. To justify the application of new or repurposed drugs in humans, pre-clinical in vivo animal models are mandatory to investigate relevant PK and PD properties and their relationship to each other. In this study, an adapted mouse model was chosen to demonstrate its suitability to provide sufficient information on the model substances GS-441 524 and HCQ regarding plasma concentration and distribution into relevant tissues a prerequisite for treatment effectiveness.

**Methods:** GS-441 524 and HCQ were administered intravenously as a single injection to male mice. Blood and organ samples were taken at several time points and drug concentrations were quantified in plasma and tissue homogenates by two liquid chromatography/tandem mass spectrometry methods. In vitro experiments were conducted to investigate the degradation of remdesivir in human plasma and blood. All pharmacokinetic analyses were performed with R Studio using non-compartmental analysis.

**Results:** High tissue to plasma ratios for GS-441 524 and HCQ were found, indicating a significant distribution into the examined tissue, except for the central nervous system and fat. For GS-441 524, measured tissue concentrations exceeded the reported in vitro EC_50_ values by more than 10-fold and in consideration of its high efficacy against feline infectious peritonitis, GS-441 524 could indeed be effective against SARS-CoV-2 in vivo. For HCQ, relatively high in vitro EC_50_ values are reported, which were not reached in all tissues. Facing its slow tissue distribution, HCQ might not lead to sufficient tissue saturation for a reliable antiviral effect.

**Conclusion:** The mouse model was able to characterise the PK and tissue distribution of both model substances and is a suitable tool to investigate early drug candidates against SARS-CoV-2. Furthermore, we could demonstrate a high tissue distribution of GS-441 524 even if not administered as the prodrug remdesivir.

## 1 Introduction

Due to threat of the current SARS-CoV-2 pandemic, scientists around the world work vigorously on the development of therapeutic options, be it new chemical entities, antibodies, or repurposed known drug molecules. In order to advance quickly from pre-clinical to clinical studies, successful *in vivo* experiments in animal models are mandatory, a fact recently acknowledged by Dinnon and colleagues [1]. An adapted mouse model could be a suitable proof-of-concept for new compounds as demonstrated for remdesivir (REM) [2]. However, to step up to humans, the relationship between pharmacodynamics (PD) and pharmacokinetics (PK) needs to be assessed in such a model to calculate an appropriate dose.

Two prominent examples of drug candidates against COVID-19 are REM and hydroxychloroquine (HCQ). REM represents a prodrug developed to quickly convert to the corresponding nucleoside (GS-441 524) intracellularly or even in plasma (Figure 1). However, using GS-441 524 would be a more direct treatment strategy which does not rely on enzymatic conversion, and thus be more suitable to demonstrate the applicability of a model. GS-441 524 is the major circulating plasma metabolite of REM but only very limited data on the PK of this compound are available. Furthermore, the efficacy of REM for COVID-19 should be re-evaluated in comparison to its main metabolite (GS-441 524), which was highly effective against feline coronavirus (FCoV) causing feline infectious peritonitis (FIP) [3-5]. In fact, theoretical considerations come to the conclusion, that the prodrug REM might not be the most appropriate nucleoside strategy for the treatment of COVID-19. After all, it was specifically designed as prodrug to target the Ebola virus [6-8], while the broad and extensive multi-organ pathology of COVID-19 could actually benefit from direct application of GS-441 524.

**Figure 1.**
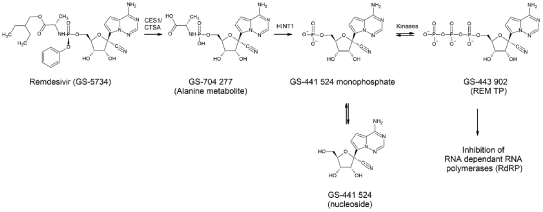
Metabolism of REM (GS-5734) leading to GS-441 524

The aim of this study was to demonstrate the applicability of mice to assess *in vivo* pharmacokinetics of two pharmacologically extremely different compounds with regards to their pharmacokinetic and physical-chemical properties. Moreover, not only plasma concentrations, but also the amount and levels in relevant tissue should be assessable to obtain information on the distribution, an important aspect regarding the question whether or not the foci of infection could even be reached, *in vivo*. In the present work, we chose a mouse model using GS-441 524 and HCQ as model compounds. We further investigated the degradation of remdesivir in human plasma and blood *in vitro* in order to add data on the aspect of its short plasma half-life and origin of GS-441 524 after administration as the phosphoramidate prodrug.

## 2 Experimental

### 2.1 Materials

GS-441 524 was purchased from BIOSYNTH® Carbosynth (Berkshire, UK), HCQ sulphate (purity: 98.5 %), [^2^H]_5_-piperacillin, [^2^H]_4_-amoxicillin and meropenem-d_6_ from Toronto Research Chemicals (Toronto, Canada). Acetonitrile Chromasolv™ Gradient was purchased from *Riedel-de Haën* (Seelze, Germany). Human EDTA Plasma was obtained from BioIVT (West Sussex, United Kingdom) Other chemicals used were of analytical grade and from VWR International GmbH (Darmstadt, Germany). Ammonium acetate, ammonium hydroxide (25%), Sodium hydroxide solution, sodium chloride solution (0.9% m/V), ethanol, propylene glycol, PEG 400, Microvetten CB300 from Sarstedt AG & Co. KG (Nümbrecht, Germany). Ultra pure water was obtained using a Milli-Q^®^ purification system (Millipore Corporation, Bedford, MA, USA).

### 2.2 Animals

Animal care was provided in accordance with the guidelines of the EU Directive 2010/63/EU and the German Animal Welfare Agency. All experiments were approved by an ethics committee and the governmental agency (Lower Saxony State Office for Consumer Protection and Food Safety; LAVES; protocol number: AZ 20A509). All efforts were made to minimize both the suffering and the number of animals. A total of 16 naive male mice (n=8 for HCQ, n=8 for GS-441 524) at 12-21 weeks of age and 22.5-38.4 g body weight on a BDF1 (C57BL6/DBA2 hybrid) background bred and housed in the institute’s facility were used. Animals were group housed (3-4 animals per cage) on standard bedding (shredded wood) and maintained on a reversed 12-h light/12-h dark cycle (lights off at 11 a.m.). Room temperature in the mouse holding room was 22 °C ± 2 °C and relative humidity was about 60%; values were recorded during the daily animal check. Food (Altromin standard diet) and water were available ad libitum and material for nest-building (paper rolls) was provided in addition to red plastic houses as enrichment.

### 2.3 Dosing and formulation

All drugs or vehicle were administered intravenously as a single injection at a volume of 5 mL/kg via the tail vein.

#### 2.3.1 GS-441 524

For pharmacokinetic studies in mice, GS-441 524 was dissolved in sterile aqua ad injectabilia with 0.625 % Ethanol, 3.75 % propylene glycol, and 5.625 % PEG 400. The pH of the drug solution and the vehicle was adjusted to 2.88 with sodium hydroxide (NaOH) and hydrochloric acid (HCl), respectively. Mice were randomly assigned to GS-441 524 at 10 mg/kg (n=6) or the respective vehicle (n=2).

#### 2.3.2 HCQ

HCQ was dissolved in sterile aqua ad injectabilia. Isotonic saline solution served as vehicle control. Solutions were prepared just prior to injection, and the final concentrations of compounds were verified from aliquots using LC-MS/MS. A cohort of mice was randomly assigned to HCQ at 5 mg/kg (free base, n=6) or the respective vehicle (n=2).

### 2.4 Collection of blood, plasma and tissue samples

The target and actual time of blood sampling were recorded relative to the time of injection. Blood samples were collected into microvette CB300 potassium-EDTA tubes, briefly mixed by slow inversion and placed on ice immediately. Within 1 hour after collection, plasma was separated by centrifugation at 3,000 g for 10 minutes at 4 °C.

Blood samples were collected from all mice immediately prior to injection of drugs or vehicle (baseline control) from the tail vein (20-30 µL), and immediately after injection from the facial, sub-mandibular, vein (20-30 µL). The plasma or whole blood was protected from light and stored at -80 °C until analyzed by LC-MS/MS for quantification.

#### 2.4.1 GS-441 524

The animal group for testing GS-441 524 was split in half to allow further repeated blood sampling at 15, 30, 60, 120 and 240 min as follows: n=3 (and n=1 vehicle) were bled at 15 and 30 min post injection from the facial vein, and at 60 min after sacrifice by decapitation from the vena cava; the other n=3 (and n=1 vehicle) were bled at 60 and 120 min post injection from the facial vein, and at 240 min after sacrifice from the vena cava.

From GS-441 524 or respective vehicle injected mice, the following organs were taken at sacrifice 60 min post injection: cerebrum, cerebellum, lung, liver, kidneys (partly separated in cortex and medulla), spleen, heart, muscle (quadriceps). For animals sacrificed 240 min post injection, additional samples were taken as follows: stomach, intestine (separated in small intestine and colon), nasal mucosa, and a sample of the cortex separated from cerebrum.

#### 2.4.2 HCQ

The animal group for testing HCQ was also split in half to allow further blood sampling. In addition to the immediate sample, blood sampling was repeated at 60 and 180 min from the facial vein, and at 360 min after sacrifice from the vena cava in one group (n=3 plus n=1 vehicle) and at 240, 480 min and 24 hrs from the facial vein, and at 30 hrs after sacrifice from the vena cava in the other group (n=3 plus n=1 vehicle). An additional 50 µL of whole blood was taken at sacrifice from the vena cava and transferred into plastic tubes without additives.

From HCQ and respective vehicle injected mice, all the above mentioned organ samples (GS-441 524) were taken, with the addition of pancreas. Surgical equipment was carefully cleaned with ddH2O and dried between samples.

In both cases, organs were quickly dried with moistened gauze to remove excess water, blood or content in case of intestines, placed in cryotubes and snap frozen in liquid nitrogen and kept at -80 °C until analyzed by LC-MS/MS for quantification.

### 2.5 Determination of plasma and tissue concentrations

GS-441 524 and HCQ concentrations were quantified in mouse plasma and several mouse tissue homogenates by two liquid chromatography/tandem mass spectrometry (LC-MS/MS) methods using an SCIEX API 6500™ triple quadrupole (GS-441 524) and an API 5500 triple quadrupole (HCQ) mass spectrometer (SCIEX, Concord, Ontario, Canada). Both instruments were equipped with turbo ion spray interface (SCIEX, Concord, Ontario, Canada). Details about the methods including quality data are described in two separate publications currently in preparation.

### 2.6 Pharmacokinetic analysis

Data acquisition and processing of raw data was performed using Analyst software version 1.7 (GS-441 524) or 1.6.2 (HCQ) (SCIEX, Concord, Ontario, Canada). Raw data were collected and processed using Microsoft Excel 2016 Version 16.0 (Microsoft Corporation, Redmond, WA, USA). Statistical calculations included descriptive analysis, simulations and visualization of results was performed with R Studio Version 1.2.5042 (RStudio Incorporation, Boston, MA, USA) running R version 3.6.3 (R Foundation for Statistical Computing, Vienna, Austria, 2020) using the packages ‘ggplot2’, ‘NonCompart’, and ‘tidyverse’ [9-12]. AUC was calculated using the linear up, logarithmic down method. The terminal slope (λ_z_) was calculated by log-linear regression of the last three data points. Published PK-data was imported using the WebPlotDigitizer version 4.3 [13].

### 2.7 In vitro degradation of remdesivir in human plasma

#### 2.7.1 Samples

Commercially available EDTA-Plasma was spiked with remdesivir at a concentration of 14 μg/mL. 3 mL Aliquots of this solution were stored for 23 hours and 40 minutes at 4 °C, at room temperature, and at 38 °C.

For high resolution mass spectrometry, 25 μL of each sample were extracted with 150 μL of acetonitrile and centrifuged for 5 min at 3 °C/14k rpm (A). 25 μL of the supernatant were diluted with 435 μL water (B). Alternatively, 100 μL of sample where extracted with 400 μL methanol (C).

For LC-MS/MS, 50 µL of each sample were combined with 300 µL internal standard solution (acetonitrile containing 50 and 250 ng/mL [^2^H]_5_-piperacillin and [^2^H]_4_-amoxicillin as internal standards) and vortex mixed for 10 seconds. After centrifugation for 3 min at 4 °C/11k rpm, 300 µL of supernatant were mixed with 300 µL of water and 1000 µL of dichloromethane and mixed thoroughly. After centrifugation for 5 min at 4 °C/3.6k rpm, the aqueous layer was transferred to an HPLC vial.

#### 2.7.2 High resolution LC-MS negative

A Sciex X500 qTOF mass spectrometer (Sciex, Darmstadt, Germany) was coupled to an Agilent Infinity 1290 UHPLC (Agilent, Waldbronn, Germany) and operated in ESI negative mode. Chromatographic separation was achieved on a Hypercarb 5 µm 100×2.1mm column (ThermoFischer Scientific, Germany) as stationary phase using 100 mM ammonium acetate at pH 9.5 (NH_4_OH 25%) and acetonitrile containing 0.1% (V/V) NH_4_OH 25% as mobile phase A and B, respectively. The gradient was programmed as follows: 0-0.5 min: 100% A, 0.5-3 min: 100→30% A, 3-3.5 min: 30→5% A, 3.5-3.6 min: 5→100%A, 3.6-5 min: 100% A. The flow rate was set to 600 µL/min.

#### 2.7.3 High resolution LC-MS positive

A Sciex X500 qTOF mass spectrometer was coupled to an Agilent Infinity 1290 UHPLC and operated in ESI positive mode. Chromatographic separation was achieved on a Kinetex F5 2.6 µm 50×4.6mm column (ThermoFischer, Germany) as stationary phase using water containing 0.1% (V/V) formic acid and acetonitrile containing 0.1% (V/V) formic acid as mobile phase A and B, respectively. The gradient was programmed as follows: 0-0.5 min: 90% A, 0.5-3 min: 90→30% A, 3-3.5 min: 30→5% A, 3.5-3.6 min: 5→90%A, 3.6-5 min: 90% A. The flow rate was set to 1000 µL/min.

#### 2.7.4 Low resolution LC-MS/MS

A Sciex 5500 triple quadrupole mass spectrometer was coupled to an Agilent 1200 HPLC and operated in ESI positive mode using the MRM experiment. Chromatographic separation was achieved on a Kinetex F5 2.6 µm 50×4.6mm column (Phenomenex, Aschaffenburg, Germany) as stationary phase using water containing 0.1% (V/V) formic acid and acetonitrile containing 0.1% (V/V) formic acid as mobile phase A and B, respectively. The gradient was programmed as follows: 0-0.3 min: 95% A, 0.3-0.7 min: 95→5% A, 0.7-1.6 min: 5% A, 1.6-1.61 min: 5→95%A, 1.61-2.5 min: 95% A. The flow rate was set to 900 µL/min from 0 to 0.3 min, changed to 1200 µL/min from 0.3 to 0.7 min and was kept at this rate until 2.5 min. MRM transitions were as follows: remdesivir: 603.3→402.0, GS-441 524: 292.2→163.1, alanine metabolite: 443.1→202.1.

## 3 Results

### 3.1 Safety

No obvious adverse effects were observed after injection of GS-441 524 and HCQ, respectively.

### 3.2 Plasma and blood concentrations

#### 3.2.1 GS-441 524

The pooled data of 6 mice was used to estimate the PK properties of GS-441 524 in wildtype mice (see Figure 2 A). Based on the estimated λ_z_ and AUC_0-∞_, the volume of distribution (V_z_) was estimated at 1.39 L/kg, whereas Clearance was estimated at 1.19 L/h/kg.

**Figure 2.**
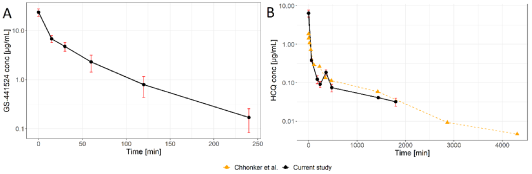
Pharmacokinetic plasma concentration vs. time profile of A: GS-441 524 in mice at 5 mg/kg (n = 6) every mouse contributed 4 measurements. means±SD B: HCQ in mice at 5 mg/kg (n = 6) each group of three mice contributed 4 and 5 measurements, respectively. means±SD overlaid with results obtained by Chhonker et al. at 5 mg/kg (dashed line with triangles) in mouse blood.

#### 3.2.2 HCQ

The pooled data of 6 mice was used to estimate PK parameters of HCQ in wildtype mice. V_z_ and Clearance were estimated at 20.1 L/kg and 0.77 L/h/kg, respectively. The mean blood-to-plasma ratio (B/P) 6 and 30 hours after infusion of 5 mg/kg HCQ was 3.13±1.43 (n = 3) and 1.31±0.104 (n = 3), respectively. The obtained plasma concentration-time profile is presented in Figure 2 B.

### 3.3 Tissue concentrations

#### 3.3.1 GS-441 524

One hour after administration of 10 mg/kg, mean tissue-to-plasma ratio (T/P) ranged from 0.14 (brain) to 2.70 (adrenal cortex). Four hours after i.v. administration, mean T/P was as high as 0.22 and 13.02 in brain and adrenal cortex, respectively (see Figure 3 A). Highest mean T/P were observed for liver (13.67), kidneys (8.61-27.04) and intestines (> 8.0). Obtained absolute tissue concentrations are presented in Table 1. The full summary is presented in Table S1.

**Table 1.** Mean observed concentrations and standard deviations (n=3) in mouse tissue following a single dose (i.v.) of 10 mg/kg of GS-441 524 or 5 mg/kg HCQ, respectively

| organ | GS-441 524 |  |  |  | HCQ |  |
| --- | --- | --- | --- | --- | --- | --- |
|  | 60 min post dose |  | 240 min post dose |  | 360 min post dose |  |
| | Conc [$\mu\text{mol/kg}$] | SD [$\mu\text{mol/kg}$] | Conc [$\mu\text{mol/kg}$] | SD [$\mu\text{mol/kg}$] | Conc [$\mu\text{mol/kg}$] | SD [$\mu\text{mol/kg}$] |
| brain | 1.24 | 0.74 | 0.13 | 0.10 | 0.29 | 0.11 |
| fat | 1.73 | 0.36 | 0.33 | 0.33 | 1.09 | 0.20 |
| cortex | - | - | 0.12 | 0.21 | 0.37 | 0.16 |
| cerebellum | 1.91 | 1.71 | 0.17 | 0.12 | 0.25 | 0.08 |
| lungs | 13.81 | 2.50 | 0.94 | 0.27 | 18.01 | 7.22 |
| heart | 14.29 | 2.99 | 1.35 | 0.52 | 5.30 | 1.78 |
| muscle | 14.57 | 3.03 | 1.82 | 1.43 | 3.47 | 0.43 |
| nose mucosa | - | - | 0.75 | 1.30 | - | - |
| stomach mucosa | - | - | 1.52 | 0.48 | 10.59 | 7.19 |
| spleen | 16.09 | 4.05 | 2.01 | 0.85 | 9.95 | 1.86 |
| stomach wall | - | - | 3.12 | 2.07 | 5.15 | 0.34 |
| small intestine mucosa | - | - | 4.69 | 2.15 | 10.92 | 7.35 |
| small intestine total | - | - | 4.37 | 1.73 | 7.26 | 4.89 |
| liver | 19.79 | 0.93 | 6.76 | 1.19 | 12.15 | 6.85 |
| left kidney | 25.67 | 7.40 | 3.99 | 3.35 | 4.88 | 1.24 |
| right kidney | 18.22 | 4.91 | 15.32 | 16.66 | 11.23 | 11.91 |
| colon mucosa | - | - | 7.05 | 5.32 | 7.11 | 3.88 |
| adrenal cortex | 25.53 | 5.46 | 5.49<sup>a</sup> | 2.18 | 14.77 | 14.56 |
| colon wall | - | - | 8.63 | 6.38 | 2.63 | 1.03 |

<sup>a</sup> only data from two animals available

**Figure 3.**
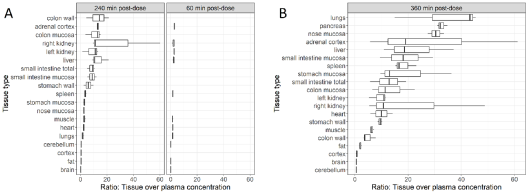
Tissue distribution in mice of A: GS-441 524 4 and 1 h post dose and B: HCQ 6 h post dose

#### 3.3.2 HCQ

Six hours after administration of 5 mg/kg, mean T/P ranged from 0.45 (cerebellum) to 34.51 (lungs). With exception of tissue found in the CNS, fat, muscle, colon and stomach wall, mean T/P was at least 10 for all tissues (see also Figure 3 B). Obtained absolute tissue concentrations are presented in Table 1. The full summary is presented in Table S2.

### 3.4 In vitro degradation of remdesivir in human plasma

TOF-MS and TOF-MS/MS experiments revealed a rapid decrease of REM at elevated temperature in plasma in vitro. At the same time, the peak due to the alanine metabolite appeared and increased accordingly. GS-441 524 was not detected in those experiments (Fig. S1-S3).

In the low resolution LC-MS/MS experiments, GS-441 524 and the alanine metabolite were present at about the same level, expressed as peak area ratio (analyte/internal standard) prior to incubation. After incubation at 38 °C the peak ratios increased by a factor of 94.3 and 29.9 for the alanine metabolite and GS-441 524 respectively. More than 99% of REM were degraded within 24 hours at 38 °C.

## 4 Discussion

### 4.1 Plasma pharmacokinetics

#### 4.1.1 GS-441 524

GS-441 524 is the major circulating plasma metabolite of REM but only very limited data is available on the PK of this compound. Most on the information obtainable from published sources refers to the PK after administration as the prodrug REM (GS-5734) either in animal models [6, 8] or healthy human subjects [14]. There is also a case report of two COVID-19 patients receiving REM [15]. Further hints on the PK of GS-441 524 can be found in the assessment report for Gilead Sciences’ Veklury (REM) [16]. To our knowledge, the only other publications investigating the PK after administration in the form of the nucleoside are investigations by Murphy and colleagues in cats [5].

In comparison to the estimated pharmacokinetics in cats, our results are in the same order of magnitude however suggesting increased clearance and reduced volume of distribution (see Table 2). These discrepancies could be due to interspecies differences, further highlighting the importance of thorough pharmacokinetic studies in species used to model COVID-19.

**Table 2.** Pharmacokinetic properties of GS-441 524 in different species

| Species | Matrix | Dose | C<sub>0</sub> [µg/mL] | C<sub>max</sub> [µg/mL] | t<sub>1/2</sub> (h) | AUC<sub>0-∞</sub> (h·µg/mL) | V<sub>z</sub> (L/kg) | CL (L/h/kg) | Ref. |
| --- | --- | --- | --- | --- | --- | --- | --- | --- | --- |
| Mouse | plasma | 10 mg/kg i.v. | 23.4 | - | 0.81 | 8.855 | 1.39 | 1.19 | current |
| CD-1 mice | blood | 20 mg/kg i.v.<sup>a</sup> | - | 10.433 | 5.7 | 42.426 | - | - | [34] |
| Ces1c<sup>-/-</sup> mouse | plasma | 25 mg/kg s.c.<sup>a</sup> | - | 1.774 | 4.92 | 7.901 | - | - | [6]<sup>b</sup> |
| rhesus monkey | plasma | 10 mg/kg i.v.<sup>a</sup> | - | 0.284 | 9.08 | 32.87 | - | - | [8]<sup>b</sup> |
| cat | plasma | 5 mg/kg i.v. | 5.01 | - | 3.84 | 13.39 | 2.07 | 0.37 | [5]<sup>b</sup> |
| human | plasma | 3 mg i.v. over 2h<sup>a</sup> | - | 3.20 | 12.9 | 0.0552 | - | - | [14] |
| human | plasma | 10 mg i.v. over 2h<sup>a</sup> | - | 9.40 | 22.0 | 0.264 | - | - |  |
| human | plasma | 30 mg i.v. over 2h<sup>a</sup> | - | 34.3 | 27.3 | 1.010 | - | 6.96<sup>c</sup> |  |
| human | plasma | 75 mg i.v. over 2h<sup>a</sup> | - | 85.8 | 26.9 | 2.470 | - | 7.02<sup>c</sup> |  |
| human | plasma | 150 mg i.v. over 2h<sup>a</sup> | - | 152 | 27.4 | 4.640 | - | 7.62<sup>c</sup> |  |
| human | plasma | 225 mg i.v. over 2h<sup>a</sup> | - | 257 | 30.6 | 7.350 | - | 8.16<sup>c</sup> |  |

Abbreviations: Ces1c<sup>-/-</sup>: carboxylesterase 1c knock-out, CD-1: caesarean derived-1
<sup>a</sup> administered as remdesivir (GS-5743)
<sup>b</sup> estimated from imported average value of PK data
<sup>c</sup> renal Clearance in L/h, **not** normalized to body weight

It was suggested by Yan and Muller, that REM *in vivo* is predominantly hydrolysed in plasma to yield GS-441 524 rather than being activated intracellularly [17]. However, this theory seems to contradict the increasing, dose dependant half-life of GS-441 524 after administration as REM (see Table 2). The dose dependency furthermore suggests that in this case half-life is not defined by the elimination rate constant. Instead, the liberation of GS-441 524 from tissue/organs saturated with the prodrug is suggested to be the rate limiting process at higher doses. This is underlined by the fact that renal clearance increases with increasing half-life (Table 2). The mass-balance study for REM (GS-US-399-4231) disclosed that renal clearance is in addition to hepatic extraction (via Alanine metabolite and the monophosphate) the most relevant elimination pathway and GS-441 524 is the predominant metabolite detected in urine (49%), followed by REM (10%). It is therefore not surprising that in one human patient with renal impairment receiving REM for the treatment of COVID-19, all GS-441 524 plasma concentrations were significantly elevated compared to another critically ill patient without renal impairment [15]. GS-441 524 exposure in an patient suffering from end stage renal disease receiving REM was dramatically increased compared to healthy volunteers or non-impaired patients [18].

#### 4.1.2 HCQ

Being a comparatively old drug, the PK of HCQ has been studied extensively. However, there is still limited information on its pharmacokinetics especially with regard to distribution. Reports considering the terminal half-life in plasma or blood are inconclusive and range from approx. 5 to 50 days [19, 20]. These values refer to an elimination half-life calculated after discontinuation from steady state. Due to the high degree of tissue distribution these long half-lives include redistribution from tissue and thus are considerably longer compared to a half-life calculated from a single dose or within steady state. The plasma half-life calculated from our data is comparable to the half-lives obtained in mice and humans (Table 4).

**Table 3.** Pharmacokinetic properties of remdesivir (GS-5734) in different species

| Species | Matrix | Dose | C<sub>0</sub> [µg/mL] | C<sub>max</sub> [µg/mL] | t<sub>1/2</sub> (h) | AUC<sub>0-∞</sub> (h·µg/mL) | V<sub>z</sub> (L/kg) | CL (L/h/kg) | Ref. |
| --- | --- | --- | --- | --- | --- | --- | --- | --- | --- |
| Ces1c<sup>-/-</sup> mouse | plasma | 25 mg/kg s.c. | - | 5.208 | 0.59 | 6.235 | 3.44 | 4.01 | [6]<sup>a</sup> |
| rhesus monkey | plasma | 10 mg/kg i.v. | 5.53 | - | 0.39 | 1.130 | 4.94 | 8.68 | [8]<sup>a</sup> |
| human | plasma | 3 mg i.v. over 2h | - | 0.575 | - | - | - | - | [14] |
| human | plasma | 10 mg i.v. over 2h | - | 0.221 | 0.66 | 0.230<sup>b</sup> | 45.1<sup>c</sup> | 45.3<sup>d</sup> |  |
| human | plasma | 30 mg i.v. over 2h | - | 0.694 | 0.81 | 0.774<sup>b</sup> | 48.8<sup>c</sup> | 42.0<sup>d</sup> |  |
| human | plasma | 75 mg i.v. over 2h | - | 1.630 | 0.90 | 2.000<sup>b</sup> | 56.3<sup>c</sup> | 39.7<sup>d</sup> |  |
| human | plasma | 150 mg i.v. over 2h | - | 2.280 | 0.99 | 2.980<sup>b</sup> | 73.4<sup>c</sup> | 51.8<sup>d</sup> |  |
| human | plasma | 225 mg i.v. over 2h | - | 4.420 | 1.05 | 5.270<sup>b</sup> | 66.5<sup>c</sup> | 43.1<sup>d</sup> |  |

Abbreviations: Ces1c<sup>-/-</sup>: carboxylesterase 1c knock-out<sup>a</sup> estimated from imported average value of PK data<sup>b</sup> mean value<sup>c</sup> body weight not available in publication parameter is given in L<sup>d</sup> body weight not available in publication parameter is given in L/h

**Table 4.** Pharmacokinetic properties of HCQ in different species

| Species | Matrix | Dose (HCQ base) | C<sub>0</sub><br>[µg/mL] | C<sub>max</sub><br>[µg/mL] | t<sub>1/2</sub> (h) | AUC<sub>0-∞</sub><br>(h·µg/mL) | V<sub>z</sub> (L/kg) | CL (L/h/kg) | Ref. |
| --- | --- | --- | --- | --- | --- | --- | --- | --- | --- |
| Mouse | plasma | 5 mg/kg i.v. | 6.41 | - | 18.2 | 5.17 | 20.1 | 0.77 | current |
| Mouse | blood | 5 mg/kg i.v. <sup>a</sup> | 1.75±0.33 | - | 12.7±1.1 | 5.58±0.88 | 17.0±4.3 | 0.9±0.2 | [27] |
| Human<sup>b</sup> | plasma | 310 mg p.o. | - | 0.403±0.134 | 172.3±39.0 | 34.36±20.42 | 37.4±28.2<sup>c</sup> | 0.16±0.09<sup>c</sup> | [40] |
| Human<sup>d</sup> | blood | 155/232.5/310 mg p.o. | - | - | 33.7<sup>e</sup> | - | 15.3<sup>c,e</sup> | 0.31<sup>c,e</sup> | [41] |
| Human<sup>d</sup> | plasma | 155/232.5/310 mg p.o. | - | - | 24.8<sup>e</sup> | - | 41.2<sup>c,e</sup> | 1.15<sup>c,e</sup> |  |
| Human<sup>f</sup> | blood | 155 or 310 mg p.o. | - | - | 42.4<sup>g</sup> | - | 8.63<sup>g</sup> | 0.14<sup>g</sup> | [24] |
| Human<sup>h</sup> | serum | mostly 310 mg p.o. | - | - | 60.2 | - | 67.9<sup>c,e</sup> | 0.78<sup>c,e</sup> | [42] |
| Human<sup>i</sup> | serum | mostly 310 mg p.o. | - | - | 25.0 | - | 25.7<sup>c,e</sup> | 0.71<sup>c,e</sup> |  |

<sup>a</sup> it was not explicitly described whether the applied dose referred to HCQ or HCQ sulfate.
<sup>b</sup> healthy male subjects
<sup>c</sup> apparent values (V<sub>z</sub>/F, CL/F)
<sup>d</sup> Lupus Erythematosus patients
<sup>e</sup> values not derived from non-compartmental analysis, but from population pharmacokinetic modelling (V/F, CL/F), t<sub>1/2</sub> derived from CL/F and V/F
<sup>f</sup> Patients with Rheumatoid Arthritis
<sup>g</sup> values not derived from non-compartmental analysis, but from population pharmacokinetic modelling and bioavailability study (V, CL), t<sub>1/2</sub> derived from CL and V
<sup>h</sup> Pregnant women with Rheumatic Disease
<sup>i</sup> Postpartal women with Rheumatic Disease

HCQ exhibits complex pharmacokinetics with extremely high volume of distribution ranging from approx. 600 to 6400 L for a 70 kg adult. The value obtained from our experiments falls in that range and is comparable to the V_z_ obtained in mice from blood concentrations and to plasma V_z_ measured in humans (see Table 4). The substance is subject to renal and hepatic elimination. Three major metabolites are known: bis-desethyl chloroquine, desethyl chloroquine, and monodesethyl hydroxychloroquine. The latter two are considered active metabolites [20]. HCQ is predominantly metabolized via CYP2C8, CYP2D6, and CYP3A4 [21] and approximately 26% of HCQ are eliminated renally as parent drug by filtration and most likely also tubular secretion [21, 22]. Due to the structural similarity to chloroquine, which is a MATE-1 substrate, secretion via this transporter seems probable [23].

Tett et al. were the first to report bioavailability and terminal half-life of HCQ. They described the pharmacokinetics of a single i.v. and oral dose using a four- and tri-exponential equation, respectively [22]. The estimated oral systemically available fraction ranged from 0.69 to 0.79. Later investigations by Carmichael and colleagues confirmed this finding with their estimate of 0.746 [24].

In plasma, the compound is bound to serum albumin and α_1_-acid glycoprotein (AAG), which is not unusual for alkaloids [19]. The plasma protein binding ranged from 34.1 to 58.7%, was dependant on the concentration of (AAG), and was higher (approx. 1.5-fold) for the S-enantiomer [25]. Thrombocytes and especially leukocytes have been identified as deep compartment for HCQ leading to a blood-to-plasma ratio (B/P) ranging from 5.4 to 10 [21, 25]. Due to the acidic environment and organelles in those cells, HCQ is trapped in the protonated form within those kinds of cells, but not in erythrocytes. To our knowledge, B/P of HCQ has not been studied in mice before and seems to be lower according to our results. This might be explained by the fact that the murine cellular immune system differs from the humane. Whereas human blood is particularly rich in neutrophils (50-70% of leukocytes) this cell type plays a minor role in the murine blood (10-25% of leukocytes) which is dominated by lymphocytes [26].

Chhonker et al. recently published an analytical method and information on the pharmacokinetics, including the metabolites of HCQ, in mice [27]. However, their method was not validated for mouse plasma and therefore not suitable to determine T/P or B/P of HCQ. It is noteworthy, that the blood concentrations in the terminal phase obtained by Chhonker et al. are approx. 1.3 times higher (B/P measured at 30 h post dose was 1.31) compared to the plasma concentrations obtained in our experiment.

Despite the accumulation of HCQ in blood cells, Chhonker et al. measured initially lower blood concentrations compared to our plasma concentrations at the same dose [27] (see Figure 2 B) underlining the time dependency of HCQ distribution. The initial concentration measured by Chhonker and colleagues most likely represents plasma concentration “diluted” with erythrocytes, as accumulation in leukocytes and thrombocytes had not yet taken place. Furthermore, it could be agreed, that for the purpose of TDM, whole blood concentrations might be more suitable compared to plasma concentrations, due to the lower variability in whole blood. On the other hand, the determination of pharmacokinetic parameters on the basis of whole blood concentrations might not be the best choice, since distribution and clearance processes usually only affect (unbound) drug molecules in the plasma and redistribution from blood cells might be slow.

### 4.2 Tissue distribution

#### 4.2.1 GS-441 524

In cynomolgus monkeys REM ([^14^C]GS-5734, 10 mg/kg), showed a tissue-to-plasma ratio (T/P) for testes and epididymis of about 1 to 1.5 after 4 h. A considerably lower amount of radioactivity was recovered from eyes and brain (T/P: ∼ 0.5 and ∼ 0.1, respectively) indicating towards a low permeability for the blood-brain barrier for REM or any of the downstream metabolites [8]. Furthermore, T/P < or > 1 indicate involvement of influx and efflux transporters which can play an important role regarding rate and extent of tissue distribution [28]. T/P < 1 is due to efflux processes as it is mediated especially in the brain via P-gp. As REM was found to be a substrate for P-gp, low T/P values found in the brain of cynomolgus monkeys are plausible. To our knowledge, GS-441 524 itself was not investigated as a transporter substrate but according to our data, tissue-to-plasma ratio in the brain is far below 1. A hypothetical affinity of GS-441 524 to P-gp, besides its low membrane permeability, would therefore also be conceivable.

In COVID-19 patients receiving REM, a CSF to plasma ratio of 25.7% was reported by Tempestilli et al. [15]. Furthermore, they found GS-441 524 concentrations in bronchoalveolar aspirate of 8.6 and 9.2 ng/mL on day 4 and 2.7 and 3.3 ng/mL on day 7, respectively (2 patients).

Our data revealed a significant distribution of GS-441 524 into numerous organs, such as liver and kidney with high K_T/P_ values both after one and after four hours. Hence, the phosphoramidate prodrug structure in the form of REM might not be necessary to achieve sufficiently high tissue and even intracellular concentrations. The high efficiency of GS-441 524 in treating FIP corroborates this finding [4]. Given the multi-organ pathology observed in COVID-19, successful treatment strategies require broad distribution. Lack of penetration into brain tissue after could explain limited efficacy of most therapeutics against neurological symptoms which are frequent and debilitating in severe COVID-19 patients [29].

As GS-441 524 represents the nonphosphorylated nucleoside form of REM, an active uptake process into the cell via equilibrative (ENT) or concentrative (CNT) transporters is plausible [30]. Yan and Muller also brought up the idea that, besides slow passive diffusion, GS-441 524 crosses membranes using nucleoside transporters [17]. A variety of nucleoside based antiviral drugs (e.g. Ribavirin) and their corresponding transporter proteins is presented by Pastor-Anglada et al. [31]. Due to the adenosine structure in GS-441 524, purine-preferring CNT2 is one of the most probable nucleoside transporter (K_m_ value of 8 µM for adenosine) to mediate intracellular uptake [31]. In comparison, K_m_ values for adenosine uptake by ENT1 (K_m_ 40 µM) and ENT2 (K_m_ 140 µM) are much higher [32]. Nevertheless, they should not be overlooked as several nucleoside transporters can be expressed by a single cell.

Reported in vitro EC_50_ values obtained from GS-441 524 against other coronaviridae range from 0.86 (±0.78) µM (HAE cells infected with MERS) to 0.78 µM (CRFK cells infected with FCoV) to 0.18 (±014) µM (HAE cells infected with SARS-CoV) [33]. Pruijssers and colleagues demonstrated that GS-441 524 was activated to the active GS-441 524 triphosphate (TP) following distribution into Vero E6 and Calu3 cells. However, bioactiviation of GS-441 524 was lower in Calu3 cells and higher in Vero E6 cells compared to REM. The activation in HAE cells was not investigated for GS-441 524. This differences in GS-441 524 TP formation could explain the differences in observed EC_50_ values against SARS-CoV-2 for REM vs. GS-441 524. EC_50_ values for GS-441 524 against SARS-CoV-2 infected Vero E6 and Calu3 cells ranged from 0.47 to 1.09 µM, respectively (depending on cell type and assay) [2]. In the light of the high efficacy of GS-441 524 against FIP and the similar EC_50_ values, these data support the hypothesis, that GS-441 524 could indeed be effective against SARS-CoV-2 *in vivo*. In our experiments, tissue concentrations in all measured organs except for the CNS and fat exceeded the EC_50_ by more than 10-fold after 60 min (assuming a tissue density of approx. 1 kg/L). Tissue concentrations in those organs were still above 1 µmol/kg after 240 min except for lungs and nasal mucosa (Table 1). Hence 10 mg/kg every 12 hours would potentially lead to tissue concentrations exceeding the *in vitro* EC_50_ for all tested coronaviridae in most organs during the greater part of the dosing interval. Low accumulation is expected after multiple dosing due to the short plasma half-life.

A recent work of Hu et al. [34] investigating the pharmacokinetics and tissue distribution of REM and its metabolites after intravenous administration of remdesivir (20 mg/kg) in mice showed high and prolonged tissue concentration of GS-441 524 and remdesivir as well as of the respective phosphorylated species. REM accumulated especially in liver, lungs, and kidneys in the form of GS-441 524 and the respective monophosphate (MP), explaining the prolonged liberation of GS-441 524 to plasma. Accumulation of GS-441 524 and the MP in liver could potentially contribute to the drugs hepatotoxicity. In comparison to our study, after 1 hour, GS-441 524 liver concentration was about 11-fold increased, whereas lung concentration was only about 5-fold higher, respectively. Four hours after administration of REM, GS-441 524 concentration in the lungs was equal compared to our results, whereas liver concentration was elevated approx. 6-fold. This could mean, that administration of GS-441 524 could be a safer alternative with regards to hepatotoxicity. Due to the less complex pharmacokinetics, prolonged or continuous infusion of GS-441 524 (with regular plasma concentration assessment) could be a more flexible and steerable therapy. Additionally, GS-441 524 exhibits improved solubility compared to REM and might not require high concentrations of sulfobutylether-β-cyclodextrin in an intravenous formulation further reducing the risk for hepatotoxicity observed in animal models [35] and nephrotoxicity. However, it should be noted, that Hu et al. did not use carboxylesterase 1c (Ces1c) deficient mice. This means, that plasma degradation of REM was significantly faster compared to humans, indicating, that accumulation in liver tissue could have been underestimated.

Furthermore, the results of Hu et al. regarding intracellular GS-441 524 and remdesivir TP concentrations revealed, that REM accounts only for a very small part in lung tissue and the major species is GS-441 524. While TP proportions increased, the percentage of intracellular GS-441 524 decreased indicating the conversion of the nucleoside into the TP [34].

#### 4.2.2 HCQ

In 1983 McChesney described the distribution of chloroquine and HCQ in albino rats in a qualitative and quantitative way. They found organ concentrations in the following order: bone, fat, and brain < muscle < eye < heart < kidney< liver < lung < spleen < adrenal tissue [36]. It was also demonstrated, that tissue accumulation did not reach steady state before seven months into treating albino rats with 40 mg/kg HCQ per day orally (stomach tube). After 30 weeks of treating albino rats with 7.8 mg/kg per day orally (stomach tube), T/P ranged from 83 (muscle) to 1122 (spleen) with the values for heart, kidney, liver, and lung in between (566, 700, 811, 883) [37].

We could confirm the extensive tissue distribution of HCQ, explaining the frequently reported high V_z_. In general, a high T/P would be desirable for an antiviral drug, since penetration into tissue is a necessary step for the intracellular uptake. Accumulation in internal organs like liver, kidney, lungs, and spleen was also observed in our experiments. The disparity between our T/P and the reported values in albino rats could be mostly contributed to the fact, that our experiment was based on a single dose and steady state pharmacokinetics have not been attained. Parts of the central nervous system did not show accumulation of HCQ, which is in accordance with earlier investigations [36] and may limit its use for the neurological manifestations potentially caused by SARS-CoV-2 penetration into the brain [29].

Reported in vitro EC_50_ values for HCQ against vero E6 infected SARS-CoV-2 cells range from 0.72 to 4.51-12.96 µM [38, 39]. With respect to the pronounced T/P, the comparatively high EC_50_ values (∼ 13µM) could be reached in our mouse model in some tissues (e.g. lungs and adrenal cortex). However, a multiple of the EC_50_ value might be necessary for a reliable antiviral effect. Due to the long terminal half-life and potentially slow tissue distribution (time to reach steady state in tissue!) sufficiently high concentrations in tissue might not be achieved in time. This could be one reason for the failure of the 5-day HCQ therapies tested [39]. A quicker tissue saturation is not feasible due to the comparatively high toxicity of HCQ. A prophylactic effect of HCQ could be studied infecting the current mouse model after long term (1-3 weeks) treatment with HCQ (5mg/kg per day) in order to reach high HCQ tissue concentration at the time of infection.

### 4.3 In vitro degradation of remdesivir in human plasma

It could be demonstrated, that REM quickly degrades to two major products in plasma *in vitro*. One of those degradation products is GS-441 524, the other is the alanine metabolite as an intermediate. GS-441 524 could only be found in the more sensitive LC-MS/MS approach. Due to the lack of external standards for the alanine metabolite, we cannot make a statement regarding the sensitivity in LC-MS/MS (the transition could not be optimized) and therefore not give a measured concentration. However, using a mass balance approach assuming that no other degradation products were formed, only a minor percentage (approx. 2.3%) of remdesivir was converted to GS-441 524 in plasma. If this finding translates to *in vivo*, the major part of plasma GS-441 524 originates from redistribution after cellular uptake of remdesivir and intracellular metabolism rather than hydrolysis in plasma as suggested by Yan and Muller (vide supra).

## 5 Conclusion

We could demonstrate that the mouse model platform is suitable to characterise the PK and tissue distribution of two extremely different compounds. We could demonstrate that GS-441 524 is distributed into tissue even if not administered as the prodrug REM. Thus, we believe that the mouse model and general procedure presented here is a useful tool in the early investigation of drug candidates targeted against SARS-CoV-2.

## Ethics approval

All experiments were approved by an ethics committee and the governmental agency (Lower Saxony State Office for Consumer Protection and Food Safety; LAVES; protocol number: AZ 20A509).

## Acknowledgements

We are grateful to Martina Gramer and Larsen Kirchhoff for technical assistance.

