## Supplemental Material for "Tissue Level Profiling of SARS-CoV-2 antivirals in mice to predict their effects: comparing Remdesivir’s active metabolite GS-441 524 vs. the clinically failed Hydroxychloroquine"

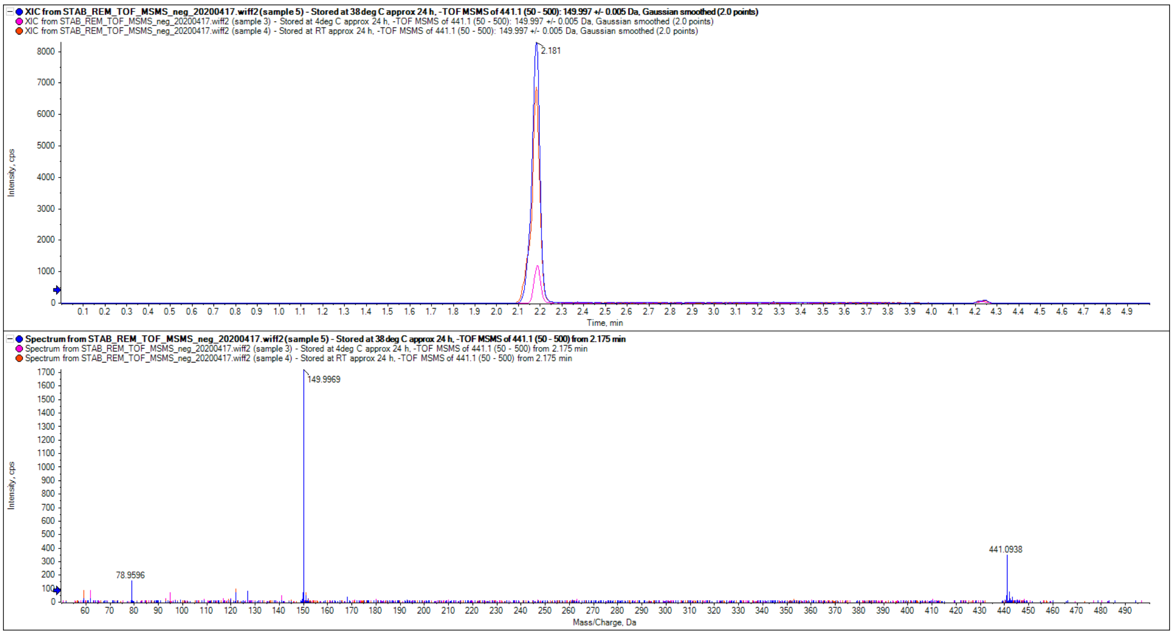


**Figure S1.** TOF-MS/MS negative mode of diluted acetonitrile preparation (B) showing the extracted ion chromatogram and the fragment ion spectrum of m/z 441.0929 ($[M-H]^-$ of the alanine metabolite) after incubation at different conditions (pink trace: 4 °C, red trace at RT, blue trace at 38 °C)

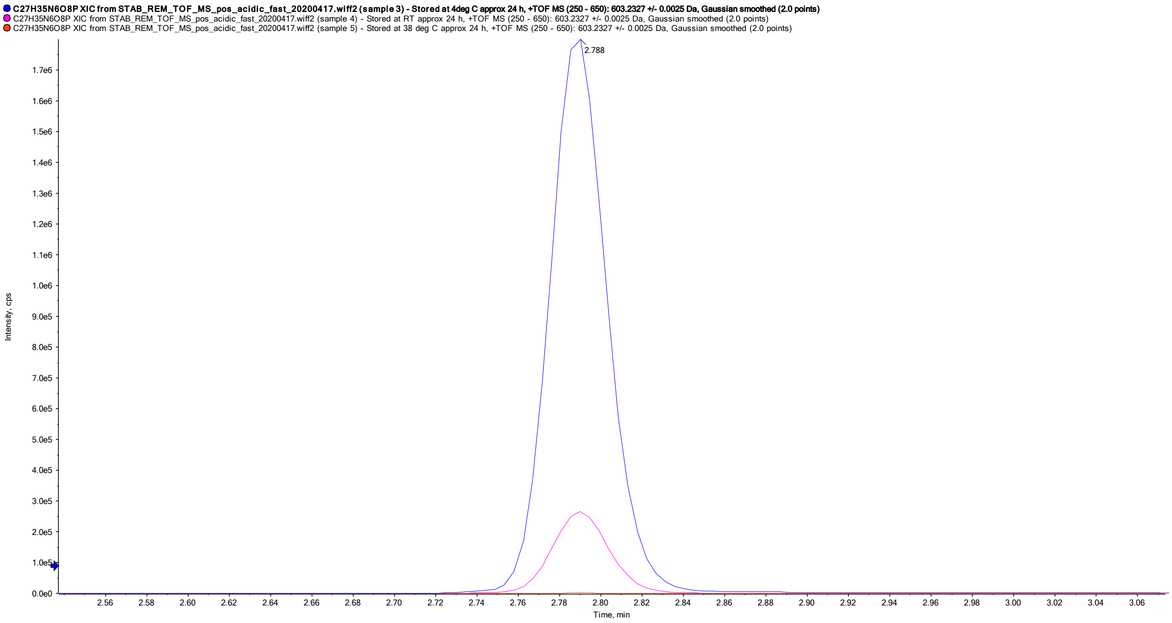


**Figure S2.** TOF-MS positive mode of methanol preparation (C) showing the extracted ion chromatogram of m/z 603.2327 ($[M+H]^+$ of remdesivir) after incubation at different conditions (red trace: 38 °C, pink trace at RT, blue trace at 4 °C)

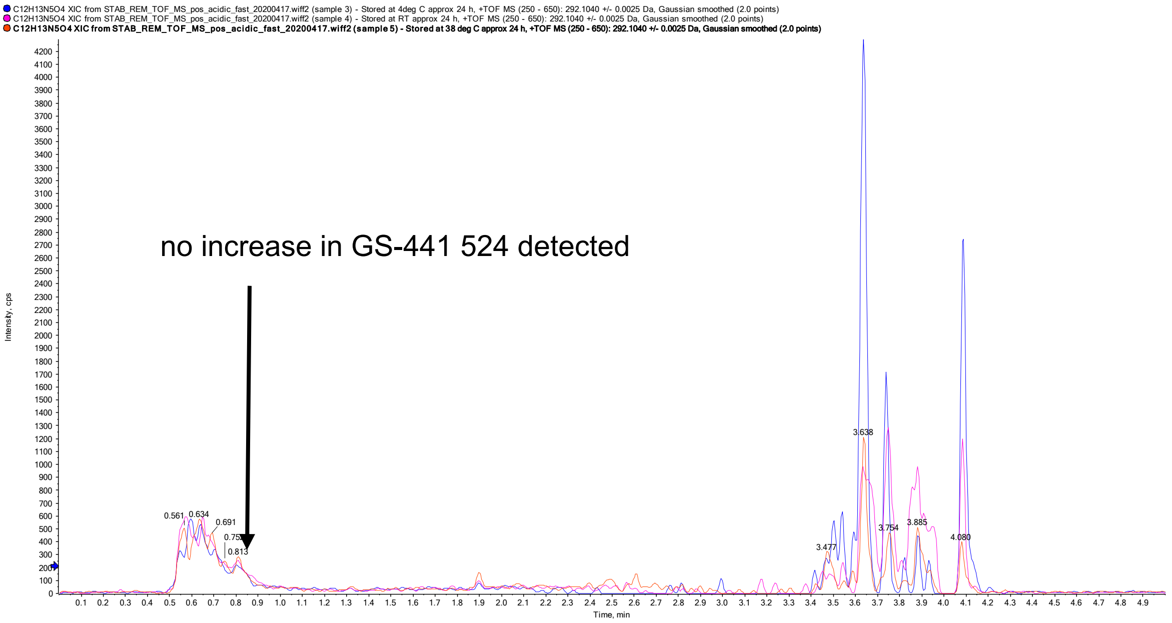


**Figure S3.** TOF-MS positive mode of methanol preparation (C) showing the extracted ion chromatogram of m/z 292.1040 ($[M+H]^+$ of GS-441 524) after incubation at different conditions (red trace: 38 °C, pink trace at RT, blue trace at 4 °C)

**Table S1.** Raw data of GS-441 524 tissue-to-plasma ratio in wild type mice

| Tissue | Time (h) | mean ratio tissue / plasma (T/P) | mean SD (T/P) | CV% |
|---|---|---|---|---|
| **Cortex[a]** | 4 | 0.41 | - | - |
| **Colon mucosa** | 4 | 10.65 | 6.05 | 56.8 |
| **Colon wall** | 4 | 13.29 | 8.54 | 64.3 |
| **Small intestine mucosa** | 4 | 8.72 | 3.40 | 39.0 |
| **Small intestines total** | 4 | 8.28 | 3.00 | 36.2 |
| **Fat** | 1 | 0.18 | 0.02 | 12.9 |
| | 4 | 0.48 | 0.28 | 54.5 |
| **Brain** | 1 | 0.14 | 0.11 | 76.0 |
| | 4 | 0.22 | 0.08 | 34.8 |
| **Heart** | 1 | 1.52 | 0.12 | 8.2 |
| | 4 | 2.46 | 0.43 | 17.3 |
| **Cerebellum** | 1 | 0.22 | 0.24 | 108.7 |
| | 4 | 0.39 | 0.30 | 75.8 |
| **Liver** | 1 | 2.16 | 0.49 | 22.9 |
| | 4 | 13.67 | 6.59 | 48.2 |
| **Lungs** | 1 | 1.47 | 0.17 | 11.3 |
| | 4 | 1.81 | 0.65 | 35.8 |
| **Stomach mucosa** | 4 | 2.83 | 0.64 | 22.5 |
| **Stomach wall** | 4 | 6.01 | 3.60 | 59.9 |
| **Spleen** | 1 | 1.69 | 0.11 | 6.6 |
| | 4 | 3.63 | 0.62 | 17.1 |
| **Muscle** | 1 | 1.55 | 0.27 | 17.1 |
| | 4 | 2.90 | 0.88 | 30.2 |
| **Nasal mucosa[a]** | 4 | 2.57 | - | - |
| **right kidney** | 1 | 2.09 | 0.96 | 46.0 |
| | 4 | 27.04 | 28.85 | 106.7 |
| **left kidney (rest)** | 1 | 2.68 | 0.04 | 1.3 |
| | 4 | 8.61 | 6.48 | 75.3 |
| **adrenal cortex left** | 1 | 2.70 | 0.23 | 8.5 |
| | 4 | 13.02 | 0.92 | 7.1 |

[a] only detectable in a single sample

**Table S2.** Raw data of HCQ tissue-to-plasma ratio in wild type mice

| Tissue | Time (h) | mean ratio tissue / plasma (T/P) | mean SD (T/P) | CV% |
|---|---|---|---|---|
| **Cortex** | 6 | 0.67 | 0.23 | 34.8 |
| **Colon mucosa** | 6 | 13.49 | 8.20 | 60.8 |
| **Colon wall** | 6 | 4.96 | 2.47 | 49.8 |
| **Small intestine mucosa** | 6 | 18.96 | 9.94 | 52.5 |
| **Small intestines total** | 6 | 12.64 | 6.76 | 53.5 |
| **Fat** | 6 | 2.02 | 0.50 | 24.6 |
| **Brain** | 6 | 0.52 | 0.16 | 31.3 |
| **Heart** | 6 | 10.00 | 4.18 | 41.8 |
| **Cerebellum** | 6 | 0.45 | 0.10 | 22.7 |
| **Liver** | 6 | 22.22 | 13.48 | 60.7 |
| **Lungs** | 6 | 34.51 | 16.92 | 49.0 |
| **Stomach mucosa** | 6 | 19.67 | 14.57 | 74.0 |
| **Stomach wall** | 6 | 9.46 | 0.88 | 9.3 |
| **Spleen** | 6 | 18.32 | 4.19 | 22.8 |
| **Muscle** | 6 | 6.37 | 0.73 | 11.4 |
| **Nasal mucosa** | 6 | 30.44 | 4.25 | 14.0 |
| **Right kidney** | 6 | 21.63 | 23.63 | 109.2 |
| **Left kidney (rest)** | 6 | 20.88 | 21.85 | 104.6 |
| **Adrenal cortex left** | 6 | 28.73 | 29.06 | 101.1 |
| **Pancreas** | 6 | 32.64 | 1.94 | 5.9 |
